# The demise of the synapse as the locus of memory: A looming paradigm shift?

**DOI:** 10.1101/082719

**Authors:** Patrick Christian Trettenbrein

## Abstract

Synaptic plasticity is widely considered to be the neurobiological basis of learning and memory by neuroscientists and researchers in adjacent fields, though diverging opinions are increasingly being recognised. From the perspective of what we might call “classical cognitive science” it has always been understood that the mind/brain is to be considered a computational-representational system. Proponents of the information-processing approach to cognitive science have long been critical of connectionist or network approaches to (neuro-)cognitive architecture, pointing to the shortcomings of the associative psychology that underlies Hebbian learning as well as to the fact that synapses are practically unfit to implement symbols. Recent work on memory has been adding fuel to the fire and current findings in neuroscience now provide first tentative neurobiological evidence for the cognitive scientists’ doubts about the synapse as the (sole) locus of memory in the brain. This paper briefly considers the history and appeal of synaptic plasticity as a memory mechanism, followed by a summary of the cognitive scientists’ objections regarding these assertions. Next, a variety of tentative neuroscientific evidence that appears to substantiate questioning the idea of the synapse as the locus of memory is presented. On this basis, a novel way of thinking about the role of synaptic plasticity in learning and memory is proposed.

## 1 Introduction

Synaptic plasticity is widely considered to provide the neurobiological basis of learning and memory by neuroscientists and researchers in adjacent fields. However, diverging opinions are increasingly being recognised (e.g., Dudai et al., 2015; Poo et al., 2016).

Within what we might call “classical cognitive science” (Piattelli-Palmarini, 2001) it has always been understood that the mind/brain is to be considered a computational-representational system. Yet, not all cognitive scientists have ever (fully) agreed with this assessment (e.g., Rumelhart et al., 1986). Actually, as of today, large parts of the field have concluded, primarily drawing on work in neuroscience, that neither symbolism nor computationalism are tenable and, as a consequence, have turned elsewhere. In contrast, classical cognitive scientists have always been critical of connectionist or network approaches to cognitive architecture (e.g., Fodor and Pylyshyn, 1988), and recent work on memory (e.g., Gallistel and King, 2009; Gallistel and Matzel, 2013; Gallistel and Balsam, 2014) has been adding fuel to the fire. Recent work in neuroscience (Johansson et al., 2014; Chen et al., 2014; Ryan et al., 2015) has now provided first tentative neurobiological evidence for the cognitive scientists’ doubts about the synapse as the locus of memory in the brain.

This paper briefly considers the history and appeal of synaptic plasticity as a memory mechanism, followed by a summary of the cognitive scientists’ objections to this idea. Next, a variety of tentative neuroscientific evidence that appears to substantiates questioning the idea of the synapse as the locus of memory is considered. On this basis, a novel way of thinking about the role of synaptic plasticity in learning and memory—mentioned only in passing in a recent commentary (Trettenbrein, 2015)—is proposed.

## 2 A tentative idea with an intuitive appeal

It was Ram´on y Cajal who first concluded from his studies of bird brains that neurons touch one another yet remain separate entities; Sherrington later coined the term “synapse” to refer to the microscopic gap between individual nerve cells (Glickstein, 2014). Subsequently, the psychologist Donald Hebb made the at first merely theoretical proposal that changes in synaptic connectivity and strength might constitute the fundamental mechanism for information storage in the brain. It it interesting to note that Ram´on y Cajal had, in a way, anticipated this conceptual move when he noted that “[…] interneuronal connectivity […] is susceptible to being influenced and modified during youthful years by education and habits” (Delgado-García, 2015, p. 6). As of today, the general idea that learning is essentially the modification of synapses in an ever-changing plastic brain (a problematic notion; see Delgado-García and Gruart, 2004; Delgado-García, 2015) has become one of the dogmas of modern neuroscience and is usually presented in popular science as well as in the scientific literature proper as an established fact and “generally accepted” (Bruel-Jungerman et al., 2007).

The fundamental principle of Hebb’s ideas on learning and memory was later poignantly summarised by Shatz as “[…] cells that fire together wire together” (1992). Hebb himself more elaborately suggested that

> [w]hen an axon of cell A is near enough to excite cell B and repeatedly or persistently takes part in firing it, some growth process or metabolic change takes place in one or both cells such that A’s efficiency, as one of the cells firing B, is increased. (Hebb as cited in Glickstein, 2014, p. 253)

While these ideas about synaptic plasticity originally were purely theoretical in nature they have long-since been confirmed experimentally with the dis¬covery of long-term potentiation (LTP) and long-term depression (LTD) as complementary neurobiological mechanisms.

In this context, it is important to point out the crucial role that (cognitive) psychology and its philosophical predecessors have played in this overall development. Not only was Hebb himself an early neuropsychologist trained by Karl Lashley, his idea that learning occurs whenever two cells fire together is clearly reminiscent of Lockean associative psychology which, in turn, can be traced back in the history of ideas all the way to Aristotle. This associative aspect of Hebbian learning has an intuitive appeal, as Gallistel and King (2009) recount tongue-in-cheek: How else could you explain that if you hear “salt” you will probably also think “pepper?” Associationism has come in different flavors since the days of Skinner, but they all share the fundamental aversion towards internally adding structure to contingencies in the world Gallistel and Matzel, 2013. Crucially, it is only against this background of association learning that LTP and LTD seem to provide a neurobiologically as well as psychologically plausible mechanism for learning and memory.

## 3 Why not (only) the synapse?

If learning actually is the association of ideas in the Lockean sense, then everything is fine and LTP and LTD provide mechanisms that enable brains to carry forward information in time. However, as the proponents of a computational-representational view of the mind/brain have been arguing almost since the inception of cognitive science against the backdrop of behav¬iorism in the 1950s, learning is not association. In similar fashion, Gallistel and collaborators (2009; 2013) have in recent years been pointing to this conceptual flaw with regard to how learning and memory are understood in neuroscience and other cognitive sciences, convincingly arguing that indeed not even the most fundamental properties of associative learning can be accounted for by LTP and LTD.

Pavlov himself already knew that the associative strength between two stimuli is determined by a multitude of experimental aspects in combination, meaning that all the different variables are encoded in a single association. In order to read information from memory it would be necessary to know about the mapping rules employed when writing that information to memory. However, in such a scenario where a variety of different variables have been mixed and encoded in a single association it is mathematically impossible to determine the value of any of the variables that entered into the original calculation: Mathematically speaking, association is a many-one function so that Pavlov’s goal of discovering general “laws of association” cannot be met. Instead, in order to regain useful information from associative strengths, it is necessary to assume that there are different mapping rules (i.e. neurobiological processes) for every synapse. It stands to reason that this is an unpleasant assumption which runs counter to Pavlov’s intent.

Furthermore, Hebbian learning rests on the idea that the neuron is the basic unit of information processing in the brain only by virtue of its connectivity profile, meaning that strength and patterns of synaptic connections constitute the way in which information is represented and stored in the brain. However, thereisno *a priori* reason why this should be the case and while the vast majority of work in cognitive neuroscience of course relies on imaging or (indirectly) recording from entire populations of neurons if not functional modules or systems, we need not and should not assume that information processing in the brain stops at the cellular level.

Based on the observation that every neuron performs a rather stereotypical computational operation on its input (Kandel et al., 2013), it seems plausible that much more information processing is actually going on inside the cell. After all, the neuron in itself is an incredibly complex and morphologically diverse structure, of which, despite all progress, we still have not reached a satisfactory understanding. Incidentally, once we adopt such a perspective we also no longer face the additional problem that changes in synaptic conductance (i.e. modification of synaptic weights) are actually not accessible to computational processes being carried out inside the cell.

### 3.1 The view from classical cognitive science

Despite the fact that the information-processing approach to the study of the mind/brain has gone somewhat out of fashion within the cognitive sciences in the past decades, Gallistel and King (2009) have presented a convincing argument for an understanding of the mind/brain as being Turing complete. (In this context, see table 1 for working definitions of key terms.) Needless to say, Gallistel and King’s original argument is by far more detailed than the sketch that I can provide here, yet the main points should surface and will hopefully suffice for further treatment.

**Table 1:** Nomenclature. This table provides brief expositions of terms and concepts from theory of computation that might not be familiar to all readers. Note that these are working definitions for the purpose of this paper, they are not meant to be exhaustive.

| Term | Exposition |
| --- | --- |
| finite-state machine | An abstract machine that can be in only one state at a time and a finite number of states in total. Its memory is defined by the number states available. |
| Turing machine | A finite-state machine extended with a so-called tape. The tape is a read/write memory component where symbols can be stored and recovered. |
| Turing completeness | Refers to the ability of a given set of instructions to simulate a Turing machine. |
| von-Neumann implementation | Denotes a common schematic circuit concept (and its many offshoots) that actually implement a universal Turing machine. |

Once we reconsider the classical cognitive scientist’s conception of the mind/brain as a computational-representational system it is evident that the brain must adhere to the abstract architectural properties of a universal Turing machine, meaning that it is capable of universal computation. This, of course, is not to say that the brain must resemble von Neumann’s implementation of a Turing machine, but that it nevertheless seems to adhere to the abstract properties of a Turing machine. Crucially, as David Marr put it, “[v]iewing our brains as information-processing devices is not demeaning […]” (2010, p. 361). It might be the case that the brain is capable of carrying out computations that a Turing machine cannot compute, but we do not know whether this is the case nor how the brain might achieve this feat.

Provided that the cognitive functions exhibited by human brains require the capabilities of Turing machines, one could quickly be led to a Scala Naturae interpretation of the evolution of computational abilities of nervous systems. Humans seem to be generalists, whereas animal learning is usually seen as highly domain-specific (e.g., Gallistel, 1999). Thus, one might be led to conclude that only human brains are capable of universal computation. In fact, some have made this exact claim with regard to human language capacities (e.g., Steedman, 2014). However, this would clearly be a mistake, as it is now understood that even insect navigation already requires the capabilities of a Turing machine (Gallistel, 1998; Gallistel and King, 2009). In other words, ”[…] ants have already climbed all the way up Natures ladder.” (Berwick and Chomsky, 2016, p. 132)

The crucial feature of a Turing machine is its memory component: the (hypothetical) machine must possess a read/write memory in order to be vastly more capable than a machine that remembers the past only by changing the state of the processor, as does, for example, a finite-state machine without read/write memory. Thus, there must be an efficient way of storing symbols in memory (i.e. writing), locating symbols in memory (i.e. addressing), and transporting symbols to the computational machinery (i.e. reading). It is exactly this problem, argue Gallistel and King (2009), that has by and large been overlooked or ignored by neuroscientists.

Now, when we are looking for a mechanism that implements a read/write memory in the nervous system, looking at synaptic strength and connectivity patterns might be misleading for many reasons. Most pressingly, as Gallistel and King point out, synapses might already be too complex in terms of implementing such a very basic function:

> In the final analysis, however, our skepticism rests most strongly on the fact that the synapse is a circuit-level structure, a structure that it takes two different neurons and a great many molecules to realize. It seems to us likely for a variety of reasons that the elementary unit in the memory mechanism will prove to be a molecular or sub-molecular structural unit. (2009, p. 282)

Hence, they suggest turning to DNA and RNA, which already implement the functionality of a read/write memory at the sub-molecular level. Interestingly, in discussing recent work on memory, Poo et al. reach a similar conclusion when they remark that “[…] some other mechanisms, potentially involving epigenomic modifications in engram neurons, appear to be necessary for memory trace storage” (2016, p. 8).

A mechanism as essential as memory has to be efficient in all respects, be it implementational complexity or energy efficiency. Another part of Gallistel and collaborators’ argument for the point of view they put forward is the observation that neural computation is demonstrably incredibly fast, therefore making it much more likely that the memory mechanism is (sub-)molecular in nature so that computational machinery and memory can be located in close physical proximity in order to minimize the distance over which a signal has to be transmitted (a process which evidently is “slow” in the nervous system in comparison to, for example, conventional computers).

### 3.2 Some tentative evidence

To this day, tentative evidence for the (classical) cognitive scientists’ reserva¬tions towards the synapse as the locus of memory in the brain has accumulated. A lot of groundbreaking work concerning the way in which the brain carries forward information in time was actually performed on comparatively simple model organisms such as *Aplysia* and has then been extrapolated to speculate about what might be going on in human mind/brains (e.g., Kandel and Siegelbaum, 2013). Interestingly, it is recent work in this exact domain which has indicated that the idea of synaptic conductance as the basic memory mechanism is insufficient and incomplete at best.

In Kandel and collaborators’ by now classic work with *Aplysia*, changes in synaptic conductivity were shown to alter how the animal reflexively responds to its environment. But not even in *Aplysia* all synapses are equally susceptible to change, many appeared not to be very plastic Kandel and Siegelbaum, 2013. Recent work with cultured *Aplysia* motor and sensory neurons by Chen et al. (2014) has revealed that long-term memories appear to persist covertly in cell bodies and can be restored after synapses have been eliminated. Long-term memory persisted after pharmacological elimination of synapses that had been produced only after learning had occurred, calling the role of synapses as the presumed locus of memory into serious doubt.

Similarly and possibly even more convincing, in a groundbreaking study bulding on earlier work (Hesslow et al., 2013) that already pointed to the mismatch between LTD in Purkinje cells and cerebellar learning, Johansson et al. (2014) investigated how the response of Purkinje cells changes during learning. Studying eyeblink conditioning, they showed that the cells could learn the temporal relationship between paired stimuli during conditioning. Strikingly and in stark contrast to widespread belief, the timing of responses exhibited by conditioned Purkinje cells after conditioning did not depend on a temporally patterned input. Consequently, Johansson et al. conclude that both, timing mechanism and memory trace, are located within the Purkinje cell itself. As they put it, “[…] the data strongly suggest that the main timing mechanism is within the Purkinje cell and that its nature is cellular rather than a network property” (Johansson et al., 2014, p. 14933).

Lastly, in a recent study supposed to demonstrate the increase in synaptic strength and density of dendritic spines during memory consoldiation, Ryan et al. (2015), to their own surprise, showed that changes in synaptic strength are not directly related to storage of new information in memory. In accordance with the literature on memory consolidation, Ryan et al. found that injection of protein synthesis inhibitors induced retrograde amnesia, meaning that the memory could not be retrieved. However, when optogenetically activating the neurons previously tagged during the conditioning process, memories could nevertheless be retrieved despite chemical blocking, indicating that the formation of synapses or strengthening of synaptic weights is not critical to memory formation as such.

### 3.3 Rate of synaptic turnover

The synaptic trace theory of memory requires synaptic conductance and connectivity to change during learning, that is when new information is being memorised. Studies by Xu et al. (2009) and Yang et al. (2009) both used two-photon microscopy, a feat of contemporary technology that makes it possible to trace individual synaptic spines over prolonged periods of time (i.e. weeks to months), in order to investigate the predicted changes in motor cortex during acquisition of a new motor skill.

As the researchers had anticipated, they found that learning of the new motor skill indeed was accompanied by the formation of new synaptic connec¬tions. Yet, the more puzzling finding of their studies is that synaptic spines were found to be still turning over at a rather high rate in absence of learning. As a matter of fact, the rate of synaptic turnover in absence of learning is actually so high that the newly formed connections (which supposedly encode the new memory) will have vanished in due time. It is worth noticing that these findings actually are to be expected when considering that synapses are made of proteins which are generally known to have a short lifetime.

Nevertheless, the observation that synapses are turning over at a high rate even in absence of learning, of course, is paradoxical. Interestingly, this was already noticed by A. von Kölliker, a contemporary of Ram´on y Cajal (Delgado-García, 2015). Today, Bizzi and Ajemian observe as an aside in a review of the current state of research on voluntary movement:

> If we believe that memories are made of patterns of synaptic connections sculpted by experience, and if we know, behaviorally, that motor memories last a lifetime, then how can we explain the fact that individual synaptic spines are constantly turning over and that aggregate synaptic strengths are constantly fluctuating? (2015, p. 91)

Just as Bizzi and Ajemian go on to describe, this finding is amongst those that are the most challenging to the idea that the synapse is the locus of memory in the brain.

Synapses have been found to be constantly turning over in all parts of cortex that have been examined using two-photon microscopy so far (see papers cited in Yang et al., 2009), meaning that (motor) memories by far outlive their supposed constituent parts. It seems that there are two possible ways of resolving this puzzle: We can either assume that memories are perpetually being retrieved from memory and re-encoded during this constant turnover, or we might conclude that the widely presumed relation between synaptic conductance and connectivity and memory is not as direct as conventional wisdom would have it. Provided that there is some merit to the idea that brain’s memory mechanism might be localized to neurons’ somata, a separation between learning and memory seems indicated.

## 4 The need to separate learning and memory

That learning and memory might be dissociated has been implicitly acknowl-edged in the neuroscience literature (e.g., Bannerman et al., 1995; Saucier and Cain, 1995). The word has not (yet) spread to other disciplines, presum¬ably because experimental results have, to a certain extent, been somewhat ambiguous Martin and Morris, 2002. Consequently, these reported tentative findings are usually not readily interpreted as evidence countering the idea of associative LTP/LTD as the putative basis for learning and memory.

A good example is spatial learning which is crucially dependent on hip-pocampus (Bannerman et al., 1995; Martin and Morris, 2002). Without going into great detail here, it can be said that the production of LTP (though not its maintenance) has been shown to crucially depend on N-methyl-D¬aspartate (NMDA) receptors located in the dentritic spine of the postsynaptic neuron (Bruel-Jungerman et al., 2007; Siegelbaum et al., 2013). It thus follows that if these receptors are chemically blocked, learning should be impaired or rendered impossible. Indeed, animals with blocked LTP exhibit impairment of their ability for spatial learning. But this general statement requires some qualifications as Bannerman et al. (1995) as well as Saucier and Cain (1995) have shown that pretraining can actually “compensate” for pharmacological blocking of NDMA receptors so that animals perform (close to) normal.

Otherwise put, when animals were pretrained in navigating in one water maze they could readily learn to navigate in a second one despite the chemical blocking of LTP. This might be interpreted as to indicate that NMDA receptors do play a role in initial learning of a new skill (i.e. navigating a maze) but do not appear to play any role in altering the specifics, or maybe better ‘contents,’ for example, when a new map is added to memory, respectively when the already existing representation is being updated. Provided that blocking of NMDA receptors did not prevent the acquisition of new information it seems reasonable to purport that a memory mechanism other than LTP was at work here, thought the nature of this mechanism remains unknown. All in all, we might take this as an indication for a dissociation of (spatial) learning and the memory mechanism(s) as such, an interpretation that has abundant representational implications (see also Gallistel and Matzel, 2013).

We might now once again turn to (classical) cognitive science and consider these findings against the backdrop that learning is highly domain-specific (e.g., Chomsky, 1975; Gallistel, 1999): An information-processing perspective on the mind/brain necessarily leads to the postulation of domain-specific learning mechanisms. Hence, based on the above-mentioned studies we might postulate that the (spatial) learning mechanism is only partially dependent on synaptic plasticity. Acquisition of the skill relies on synaptic plasticity and thus a process of neural reorganisation, whereas altering the specifics (i.e. acquisition of new information, respectively “updating” of information already stored in memory) does not. It follows that LTP and thus synaptic plasticity cannot provide the brain’s basic memory mechanism.

In the sense of Gallistel and King (2009), learning is the process of extract¬ing information from the environment, whereas memorizing is the processes of storing this information in a manner that is accessible to computation. It is interesting to note that once learning and memory are conceived of as separate processes, the above-mentioned observation that synaptic spines are still turning over at a very high rate in absence of learning does no longer pose such a severe problem. In somewhat similar fashion, we can interpret the findings of Ryan et al. (2015) against this background, so that we might say that in their study information was extracted from the environment (i.e. learning occurred) and stored in memory independently of the process of memory consolidation, that is alteration of synaptic weights and connectivity.

Lastly, all of this is not to say that synaptic plasticity and networks are of no importance for learning and memory. Fodor and Pylyshyn (1988) already reviewed the implications of connectionist models, concluding that connection¬ism might best be understood as a neutral “theory of implementation” of the actual cognitive architecture, provided that one gives up anti-representational tendencies inherent to the approach. As a consequence, the question no longer is whether symbolic representations are “real,” but how (i.e. on what level) they are actually implemented in the brain. The challenge for critics of the synaptic plasticity hypothesis will therefore be to come up with concrete suggestions for how memory might be implemented on the sub-cellular level and how cells then relate to the networks in which they are embedded.

## 5 Rethinking synaptic plasticity

The realization that the synapse is probably an ill fit when looking for a basic memory mechanism in the nervous system does not entail that synaptic plasticity should be deemed irrelevant. Quite to the contrary, there of course is ample and convincing evidence that synaptic plasticity is a prerequisite for many forms of learning (see e.g., Martin and Morris, 2002; Münte et al., 2002; Bruel-Jungerman et al., 2007; Jäncke, 2009; Dudai et al., 2015; Ryan et al., 2015; Poo et al., 2016). However, it occurs to me that we should seriously consider the possibility that the observable changes in synaptic weights and connectivity might not so much constitute the very basis of learning as they are the result of learning.

This is to say that once we accept the conjecture of Gallistel and collabo¬rators that the study of learning can and should be separated from the study of memory to a certain extent, we can reinterpret synaptic plasticity as the brain’s way of ensuring a connectivity and activity pattern that is efficient and appropriate to environmental and internal requirements within physical and developmental constraints. Consequently, synaptic plasticity might be understood as a means of regulating behavior (i.e. activity and connectivity patterns) only after learning has already occurred. In other words, synaptic weights and connections are altered after relevant information has already been extracted from the environment and stored in memory.

Over roughly the last decade, evidence that supports such an interpretation has been piling up, suggesting that the brain is (close to) “optimally wired.” It seems that axons and dendrites are close to the smallest possible length, at least within a cortical column (Chklovskii et al., 2002; Chklovskii, 2004) and possibly also globally (Cherniak et al., 2004; Lewis et al., 2012; Sporns, 2012). As noted by Chklovskii et al., this “optimality” has further pressing implications for the idea that the synapse is the locus of memory, after all,

> […] an increased number of synapses could not be accommodated without degrading performance in some way because the cortex is already optimally wired in the sense that the number of synapses is already maximal (2002, p. 345).

The role of synaptic plasticity thus changes from providing the funda¬mental memory mechanism to providing the brain’s way of ensuring that its wiring diagram enables it to operate efficiently with regard to environ¬mental and internal pressures. Viewed against the background that synapses are practically unfit to implement the cognitive scientists’ beloved symbols, it seems that we seriously have to consider that synaptic plasticity might not implement a memory mechanism as such. Instead, changes in synaptic conductance and connectivity might provide a bundle of mechanisms which regulate and ensure that the network and its modules perform and interact efficiently.

In this regard, it is vital to note that while cognitive science tells us that learning is domain-specific, these observations unfortunately cannot tell us whether the basic memory mechanism is rather uniform or not. An evolutionary argument could be put forward in favour of a view where the basic memory mechanisms is highly conserved, but such a theory has not yet been confirmed to facts. If memory actually turns out to be sub-cellular in nature, synaptic plasticity would of course not be rendered irrelevant. However, what would change is the function commonly attributed to synapses: For example, one possibility is that synapses could be understood as providing “access points” to information already stored in memory inside the cell (Ryan et al., 2015), instead of a way of carrying forward information in time. Memories stored in cells could thus possibly be considered to be synapse-specific, meaning that activating different synapses will elicit different events in the cell.

## 6 A tentative outlook

To sum up, it can be said that when it comes to answering the question of how information is carried forward in time in the brain we remain largely clueless. Fittingly, in a recent autobiographical account of his research, scientific career, and personal life, Michael Gazzaniga commented on the current problems of (cognitive) neuroscience, concluding that “[…] neuroscience still has not collected the key data because, to some extent, it is not known what that key data even is” (Gazzaniga, 2015, p. 190).

Apparently, very much as Marr (2010) envisioned, the “classical” cognitive scientist’s analysis of the information-processing problem at stake in the study of memory now has yielded first hints with regard to where neurobiologists should be looking for this key data when studying the brain’s fundamental memory mechanism(s): inside the cell. Tentative evidence from a wide variety of work in neuroscience seems to provide support for the idea that the synapse is an ill fit when looking for the brain’s basic memory mechanism: memory persists despite synapses having been destroyed and synapses are turning over at very high rates even when nothing is being learned. All things considered, the case against synaptic plasticity is convincing, but it should be emphasised that we are currently also still lacking a coherent alternative.

Adolphs (2015) optimistically listed the problem of how learning and memory work among those that he expects to be solved by neuroscientists within the next 50 years. We shall see how this turns out, but, if anything, the evidence and recent findings discussed here seem to indicate to me that we will have to rethink many of the basic propositions in the cognitive sciences and especially neuroscience in order to actually achieve this. Yet, it is not at all implausible that in the years to come we might see the paradigm shift that Gallistel and Balsam (2014) have been calling for.

